# (p)ppGpp-dependent activation of gene expression during nutrient limitation

**DOI:** 10.1101/2025.04.25.650602

**Authors:** Sathya Narayanan Nagarajan, Adam Rosenthal, Jonathan Dworkin

## Abstract

As rapidly growing bacteria begin to exhaust nutrients, their growth rate slows, ultimately leading to stasis or quiescence. Adaptation to nutrient limitation requires widespread metabolic remodeling that leads to lower cellular energy consumption. Examples of such changes include attenuated transcription of genes encoding ribosome components, in part mediated by the phosphorylated nucleotides guanosine tetra- and penta-phosphate, collectively (p)ppGpp. In addition, genes encoding proteins that facilitate survival nutrient limitation exhibit increased expression. An example is the *hpf* gene, encoding a broadly conserved protein responsible for protecting the ribosome from degradation under conditions limiting for ribosome synthesis. Here we show that (p)ppGpp plays a key role in the transcriptional activation of *hpf* as *B. subtilis* cells exit rapid growth. Specifically, we demonstrate that *hpf* transcription during nutrient limitation requires an RNA polymerase holoenzyme containing the alternative sigma factor σ^H^, encoded by *sigH*, whose expression is normally inhibited by the AbrB repressor. However, when global protein synthesis decreases, in part dependent on (p)ppGpp, AbrB levels fall, leading to increased *sigH* transcription and, consequently, *hpf* activation. This mechanism couples a key physiological consequence of nutrient limitation – reduced protein synthesis – with specific gene activation, thereby linking transcriptional and translational regulation. Finally, we demonstrate that (p)ppGpp is necessary for the gene expression underlying the elaboration of developmental fates including sporulation and genetic competence.

**Importance:** Bacteria often experience nutrient limitation and, in response, they attenuate energetically costly metabolic processes like protein synthesis. At the same time, however, they stimulate the expression of a subset of proteins that facilitate survival under this condition. This study identifies a new molecular mechanism in the model Gram-positive bacterium *Bacillus subtilis* responsible for gene expression in response to nutrient limitation that couples reduced global protein synthesis with increased transcription of specific genes. This mechanism mediates the elaboration of developmental fates including sporulation and genetic competence that are known responses to nutrient limitation in this organism.

## Introduction

Bacterial adaption to environmental changes typically includes a transcriptional response. This can be relatively narrow, as in the case of two-component signaling which usually involves a limited number of genes under the control of a single transcription factor (e.g., a response regulator). Alternatively, this can be broad, such as the widespread transcriptomic changes mediated by the nucleotide second messengers guanosine tetra- and penta-phosphate, collectively (p)ppGpp. These molecules are synthesized in response to nutrient limitation, in specific to diminished amino acid availability as reflected in the presence of uncharged tRNA molecules in the ribosome A site. Diverse bacteria including *E. coli* (1-4), *Synechococcus elongatus* (5, 6), and *Staphylococcus aureus* (7) exhibit extensive transcriptional reprogramming as a consequence of (p)ppGpp synthesis.

Although the particular physiological conditions of each of these studies differ, a consistent feature is the presence of negative transcriptional changes. In the Gram-negative *E. coli*, (p)ppGpp binds to two sites on RNA polymerase. Binding at both Site 1 and Site 2 is necessary for complete transcriptional inhibition, most likely resulting in changes to promoter kinetics (8). The protein DksA interacts with RNAP near Site 2, thereby enhancing (p)ppGpp inhibition. In Gram-positive species, the RNAP residues that (p)ppGpp contacts are not conserved and a DksA homolog has not been identified. It has been proposed that the decline in [GTP] following stringent response activation could negatively affect initiation kinetics at promoters starting with a guanosine (9, 10). However, evidence that decreases in GTP levels are not necessarily correlated with increases in (p)ppGpp levels (11), suggests that this mechanism may not be sufficient.

Although, analysis of (p)ppGpp in transcription has historically focused on its inhibitory role, some early studies identified positively regulated genes. Transcription of the *E. coli his* operon is depressed in a strain lacking *relA* (12), consistent with the effect of amino acid availability on the activation of the RelA (p)ppGpp synthase. Following this study, other amino acid biosynthetic genes were observed to be under positive control by (p)ppGpp (13). The absence of (p)ppGpp has been observed previously to prevent activation of genes that mediate the response to energy limitation. For example, many genes in the cyanobacterium *S. elongatus* induced upon exposure to dark are (p)ppGpp-dependent (5). Expression of *E. coli lrp*, encoding a global regulator of stationary phase expressed genes (14), greatly increases as cells exit growth and is dependent on (p)ppGpp. However, the mechanistic basis for activation (as contrasted with inhibition) in these contexts is unclear.

The *hpf* gene is positively regulated by (p)ppGpp in diverse species (e.g., *S. elongatus* (5), *P. aeruginosa* (15), and *B. subtilis* (16, 17)). The HPF protein is responsible for ribosome dimerization, and thereby ribosome stabilization, in post-exponential phase (18). Here we show that cells exiting exponential growth have decreased levels of the AbrB repressor that governs transcription of *sigH*, encoding the σ^H^ factor necessary for *hpf* transcription. The consequent increase in σ^H^ activity results in increased HPF expression, even in the context of global translation attenuation mediated in part by (p)ppGpp (19). We further demonstrate that additional genes under control of σ^H^ are also subject to positive control by (p)ppGpp. Specifically, we show that this mechanism is important to the patterns of gene activation during conditions of nutrient limitation when *B. subtilis* differentiates into alternative cell fates including sporulation and genetic competence.

## Results

We assayed *hpf* transcription by monitoring a strain carrying a fusion of P*_hpf_* and firefly luciferase (P*_hpf_*-luc) integrated in the chromosome. Firefly luciferase is unstable in *B. subtilis*, with a half-life ∼5 min (20), making it a useful reporter of gene expression dynamics. Expression of P*_hpf_*-luc occurred approximately as the cells stopped growing exponentially in a defined medium (S7/glucose) and entered transition phase (Fig. 1A), qualitatively similar to that observed using Northern analysis of *hpf* transcription in a rich growth medium (16). To more precisely characterize this growth transition, we computed the growth rate over the growth curve, observing a steep fall (∼2 fold) in growth rate (Fig. 1B) that coincided with a sharp increase in *hpf* transcription (P*_hpf_*-luc activity). The usual expectation is that increased transcription results in increased protein production. However, the known positive correlation between growth rate and protein synthesis (21) suggests that under conditions where growth rate is falling, predicting HPF abundance may not be straightforward.

**Figure 1.**
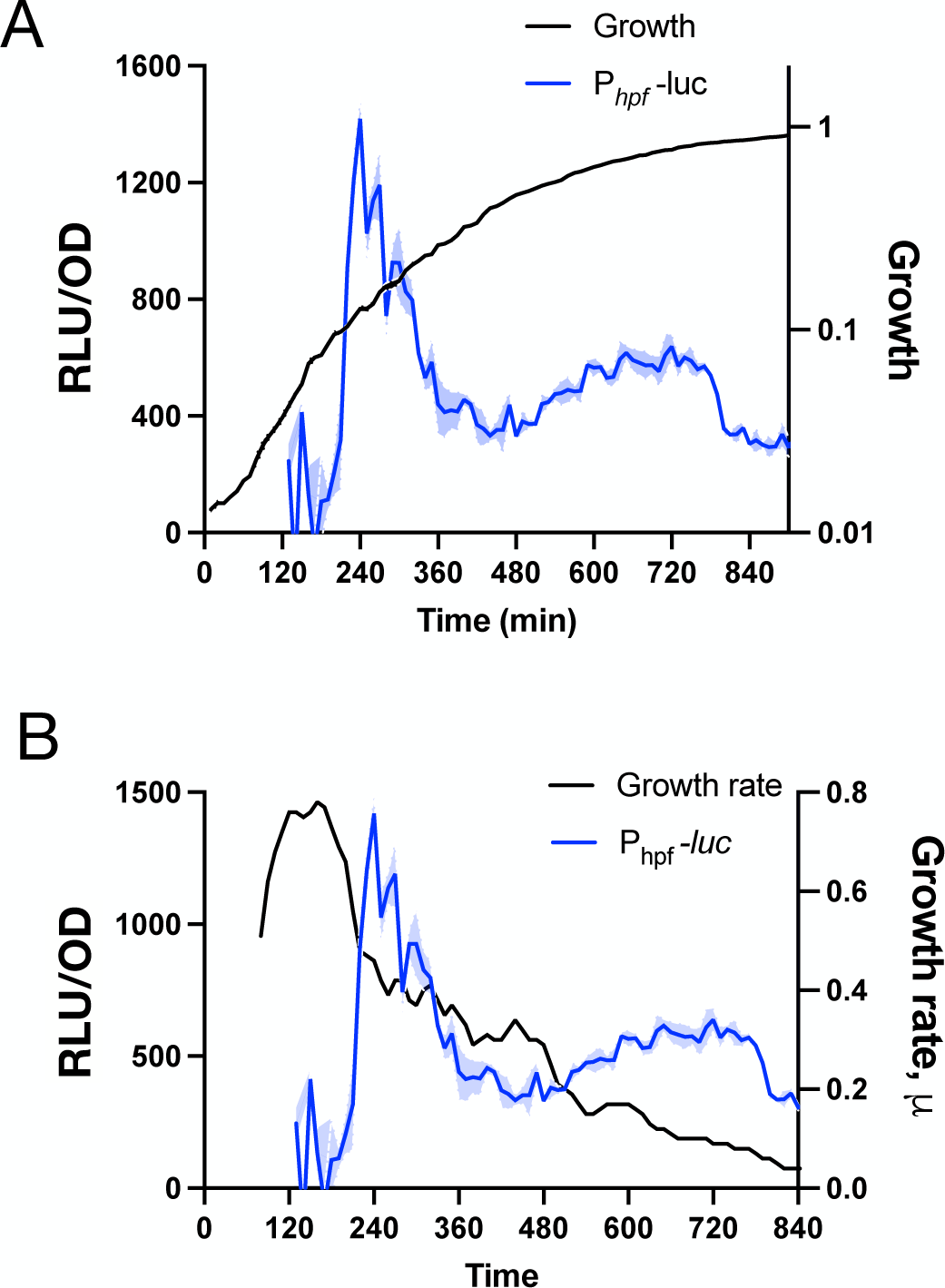
Expression of P*_hpf_*-luc and growth rate. **A,** growth (black, OD_600_) and luminescence (RLU/OD_600_; blue) of P*_hpf_*-luc expressing strain (JDB4811). **B,** luminescence (RLU/OD; blue) and growth rate (black) of P*_hpf_*-luc strain. Growth rate/hour (μ) at a given time (t) is defined as log_2_[OD_600_(t)-OD_600_ (t_-60_)].

To address this, we obtained samples of cell lysates at points in the growth curve near the spike in *hpf* transcription (Fig. 2A, arrows) and probed them with an anti-HPF antibody. This analysis revealed that HPF synthesis initiated at approximately the same point in the growth curve as the increase in luciferase activity (Fig. 2B, ‘wild-type’), similar to that observed in rich growth media (17, 22). To confirm that this interval of growth slowdown is characterized by reduced protein synthesis, we monitored incorporation of a modified puromycin (OPP) into newly synthesized polypeptide chains that can be visualized by click chemistry with a fluorophore (19). We observed that global protein synthesis decreased during the same interval ((Fig. 2C, D; 200-240 minutes) when both *hpf* transcription (Fig. 2A) and HPF protein synthesis (Fig. 2B, left) increased.

**Figure 2.**
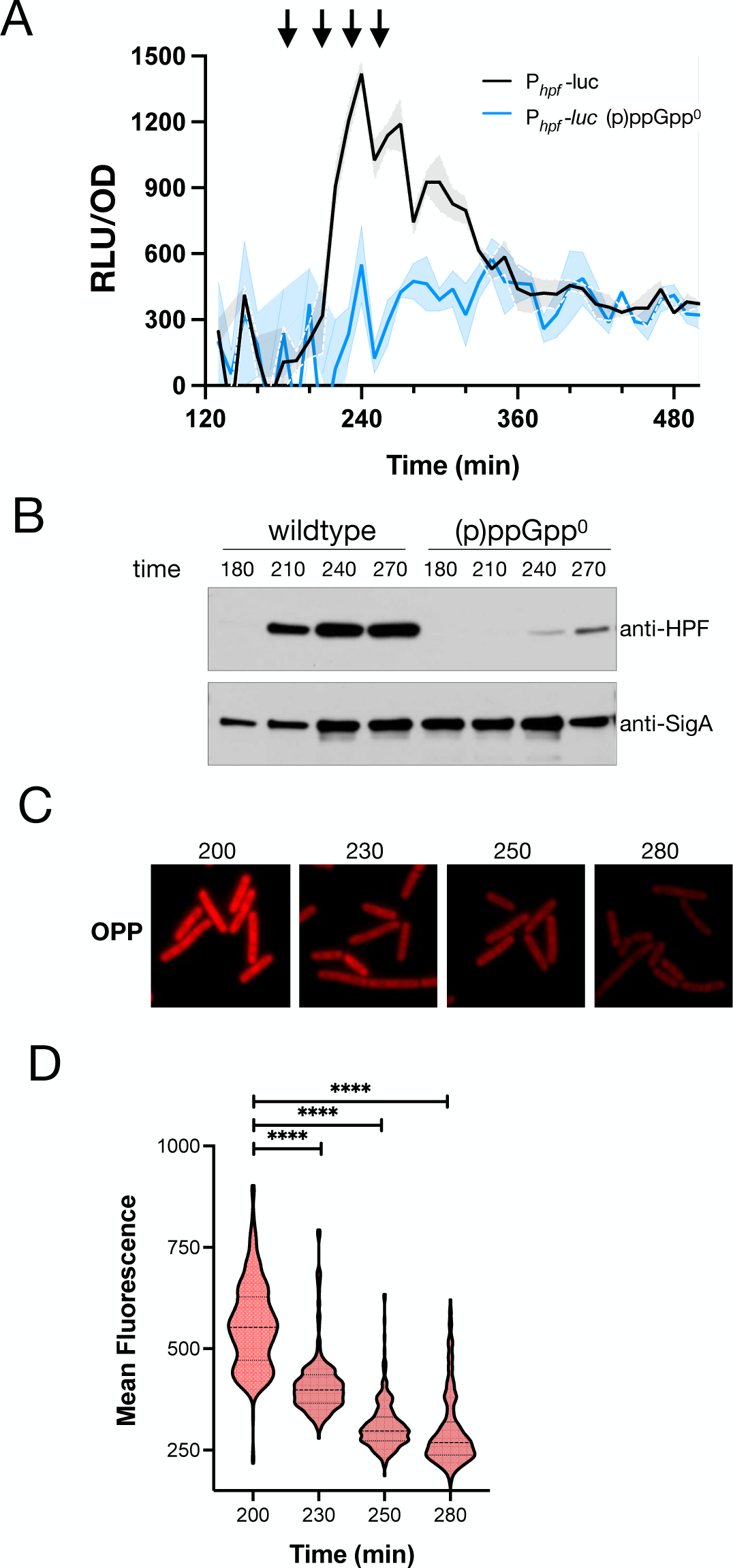
(p)ppGpp is essential for activating *hpf* transcription. **A,** luminescence (RLU/OD_600_) of wildtype (JDB4811) and (p)ppGpp^0^ (JDB4812) strains harboring P*_hpf_*-luc. Data is a representative of three independent experiments. **B,** immunoblot analysis of HPF protein (upper panel) was conducted using anti-HPF antibodies and cell-free extracts of the wildtype and (p)ppGpp^0^ strain obtained at specified time points (min). Abundance of SigA (lower panel) monitored by anti-SigA antibodies that serve as loading control. **C**, representative fluorescence microscopy images of wildtype strain labelled with OPP at specified time points (min). **D,** population distribution of mean fluorescence (in arbitrary units, AU) in the wildtype strain (JDB4811) at specified times according to C. **** indicates a two-tailed P-value, derived from a non-parametric Mann-Whitney test, of P<0.0001. Shown is a representative of three independent experiments. ∼300 cells/time point were analyzed.

These results suggest that a regulatory mechanism specifically active at this inflection point of the growth curve underlies increased *hpf* transcription. A possible component of such a mechanism is the alarmone (p)ppGpp since it regulates cellular processes in response to changes in growth (19). Consistently, the (p)ppGpp synthase Rel is necessary for *hpf* transcription in *B. subtilis* (16). However, two additional (p)ppGpp synthases – SasA and SasB - were identified subsequent to the publication of this report (23, 24). We therefore examined P*_hpf_*- luc activity in a strain lacking all three *B. subtilis* (p)ppGpp synthases ((p)ppGpp^0^), observing that it was almost completely attenuated (Fig 2A, blue). Consistently, HPF protein production was also substantially reduced in this strain (Fig. 2B, right, ‘(p)ppGpp^0^’).

To dissect the role of each of the (p)ppGpp synthases in the activation of P*_hpf_*, we monitored luciferase activity in three strains, each carrying P*_hpf_*-luc and a mutation in a single synthase. In a strain expressing a Rel mutant containing a loss-of-function mutation (Y308A) in the catalytic domain (Rel*), luciferase activity was modestly attenuated whereas *sasA* or *sasB* deletions had little effect (Fig. S1). However, mutations in (p)ppGpp synthases differentially affect protein synthesis (e.g., a Δ*sasB* strain exhibits higher global protein synthesis whereas a Δ*sasA* strain is lower compared to the wildtype parent (25)), and thus could affect luciferase activity independent of an effect on transcription. We therefore employed Fluorescence *In-Situ* Hybridization (FISH), a technique independent of protein synthesis, and investigated the effect of individual mutations on a reporter composed of P*_hpf_* fused to *gfp* (P*_hpf_*-GFP^mut2^) using oligonucleotides complementary to *gfp* mRNA. At an early transition time point, similar to P*_hpf_*- luc induction, both the *rel^Y308A^*and the Δ*sasB* mutations attenuated P*_hpf_* transcription, whereas a Δ*sasA* mutation increased it (Fig. S2). These results are consistent with the function of Rel and SasB as (p)ppGpp synthases (23, 24) and SasA being a negative regulator of SasB (25).

What accounts for the requirement of (p)ppGpp for *hpf* transcription and subsequent HPF protein production? The sigma factors σ^H^ and σ^Β^ bind defined sites in the *hpf* promoter and strains carrying deletions in their respective genes, *sigH* and *sigB*, exhibit attenuated *hpf* transcription (16). To circumvent the potential impact of deletions on overall cellular physiology, we designed P*_hpf_*-luc reporter fusions containing scrambled σ^H^ (P*_hpf-sigH_*_*_-luc) or σ^Β^ binding sites (P*_hpf-sigB_*_*_ -luc) or both (P*_hpf-sigHB*_* -luc) (Fig. 3A; Fig. S3, S4, respectively). A strain carrying the mutant P*_hpf-sigH_*_*_-luc reporter exhibited substantially attenuated luciferase activity as compared to the P*_hpf_*-luc wildtype reporter (Fig. 3B). Expression of a reporter lacking both σ^Β^ and σ^H^ binding sites (P*_hpf-sigHB*_*-luc) was even lower (Fig. S4), indicating that σ^Β^ had a residual effect on activity of the P*_hpf-sigH_*_*_-luc reporter, consistent with the defect of the P*_hpf-sigB_*_*_-luc reporter (Fig. S3). The transition state regulator CodY also binds upstream of *hpf* (26), suggesting it regulates *hpf* transcription. However, a reporter with a scrambled CodY binding site (P_hpf-CodY*_-luc) exhibited a similar pattern of activation as P*_hpf_*-luc (Fig. S5), indicating that CodY does not affect *hpf* transcription under our experimental conditions.

**Figure 3.**
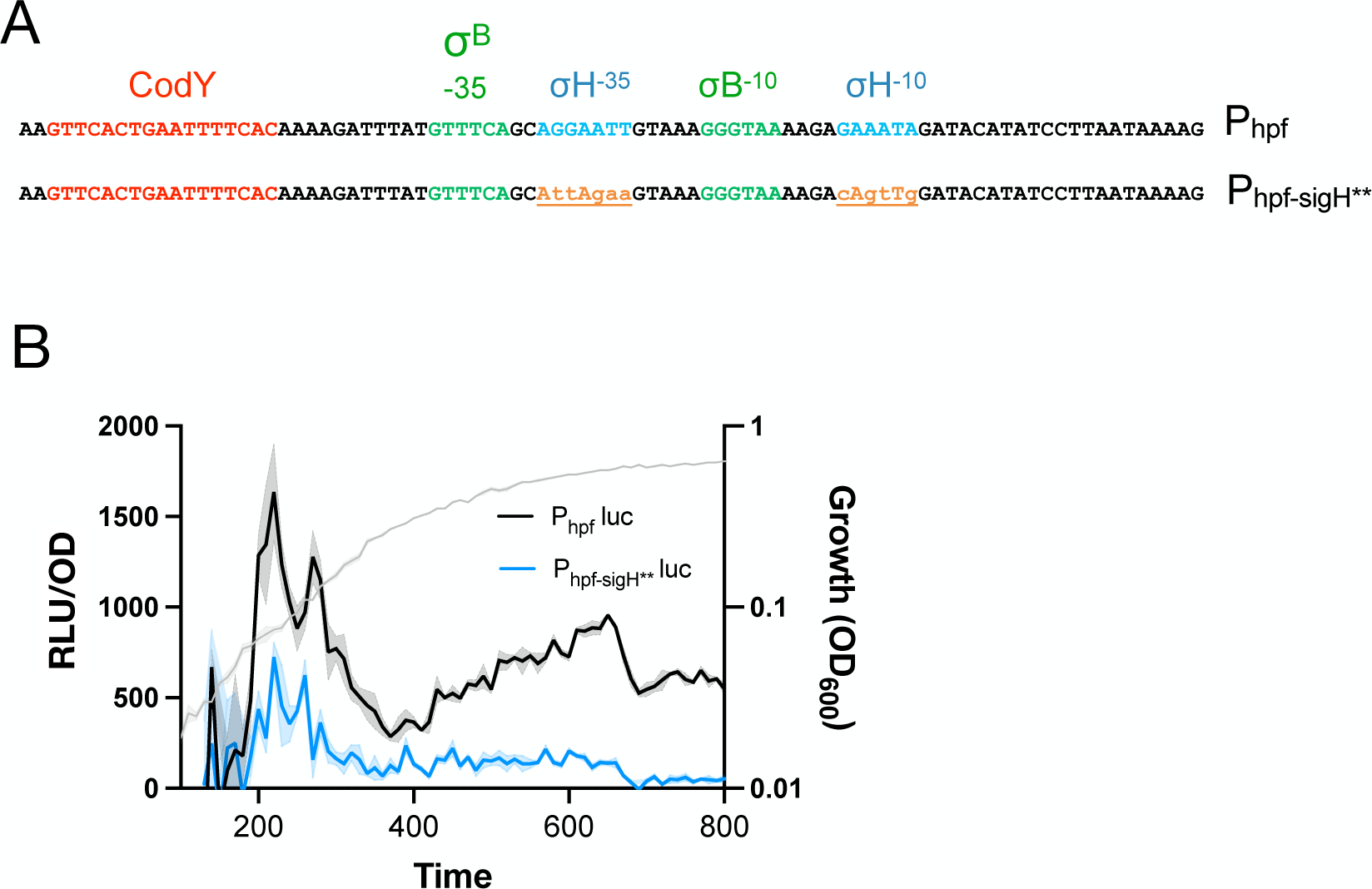
SigH is vital for P*_hpf_* activity. **A,** schematic representations of the *hpf* promoter with known binding sites of regulatory factors (top) and a mutant promoter (P*_hpf-sigH*_*) harboring mutations in the *sigH* binding site (bottom). **B**, growth (OD_600_) (gray) and luminescence of strains expressing either P*_hpf_*-luc (black, JDB4811) or P*_hpf-sigH*_*-luc (blue, JDB4813) reporters. Shown is a representative of three independent experiments.

The key role played by σ^H^ in *hpf* transcription suggests that regulation of σ^H^ expression could be the locus of control by (p)ppGpp. To examine transcriptional regulation of *sigH*, we constructed a fusion of P*_sigH_* to luciferase (P*_sigH_*-luc), which exhibited a pattern of expression similar to P*_hpf_*-luc, with a sharp peak starting at ∼180 min (Fig. 4, black). And, again similar to P*_hpf_*, expression of P*_sigH_*-luc was significantly attenuated in a (p)ppGpp^0^ strain lacking all three (p)ppGpp synthases (Fig. 4, blue), indicating that (p)ppGpp is likely affecting a regulatory step upstream of σ^H^ expression. A key regulator of *sigH* transcription is AbrB, a repressor of many transition-phase genes including *sigH* (27). We confirmed this repression occurs under our experimental conditions by examining P*_sigH_*-luc in a strain lacking *abrB*, observing that *sigH* transcription was increased as compared to the wildtype parent (Fig. S6). Thus, since decreased AbrB abundance leads to *sigH* activation (28) and consequently increased *hpf* expression, changes in AbrB levels could be responsible for the observed increases in P*_sigH_*-luc and P*_hpf_*-luc expression.

**Figure 4.**
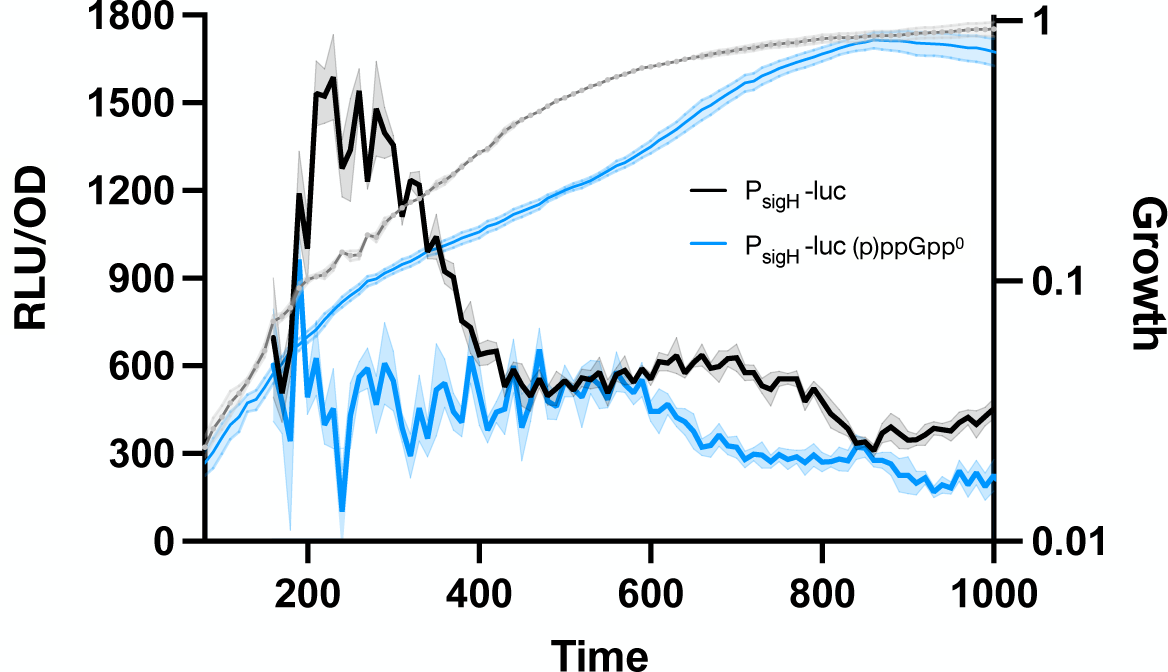
(p)ppGpp modulates *sigH* transcription. Growth (OD_600_) and luminescence (RLU/OD_600_) of P*_sigH_*-luc-containing wild-type (black, JDB4819) and (p)ppGpp^0^ strains (blue, JDB4820).

We therefore investigated AbrB abundance immediately prior to these peaks, reasoning that any changes would precede any such transcriptional changes. We initially attempted Western analysis using an anti-AbrB antibody, but were unable to consistently observe robust changes in AbrB protein levels. Alternatively, we measured the fluorescence of single cells of a strain expressing a functional AbrB-GFP fusion expressed at the endogenous locus (29). We monitored a culture expressing both *P_hpf_-*luc and AbrB-GFP and obtained samples of cells at time points before and during the initial period of the peak (Fig. 5A, arrows). Analysis of single cells by microscopy (Fig. 5B, C, ‘wildtype’) revealed a decrease in the average AbrB-GFP signal (∼40%) around the time of increase in *P_hpf_-*luc activity, consistent with our hypothesis that lower AbrB levels accompany an increase in *hpf* transcription.

**Figure 5.**
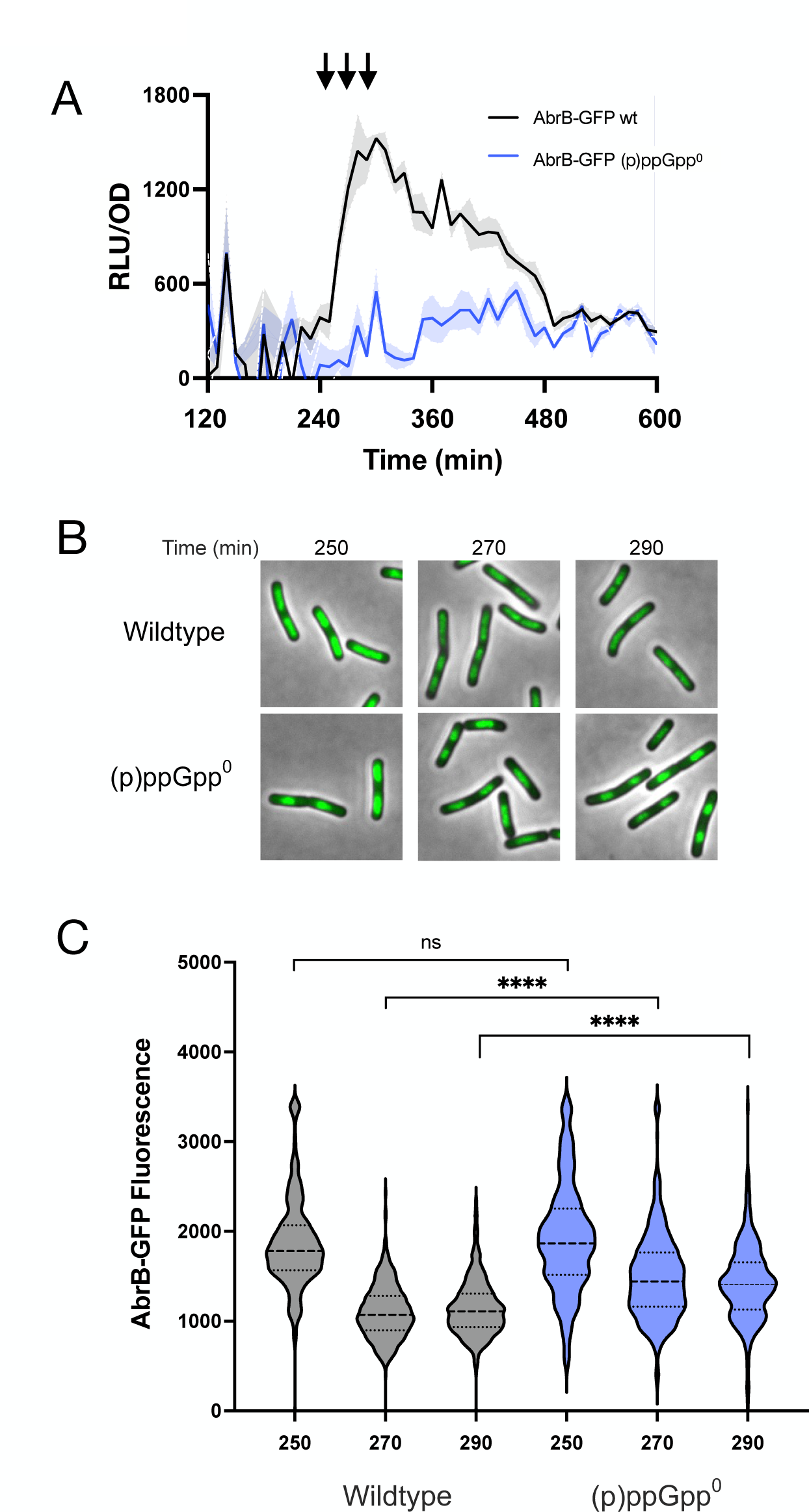
(p)ppGpp^0^ mutant has higher levels of transcriptional repressor AbrB. **A,** luminescence (RLU/OD_600_) of wild-type (black, JDB4823) and (p)ppGpp^0^ (blue, JDB4824) strains containing P*_hpf_*-luc and expressing the AbrB-GFP fusion. **B,** aliquots were obtained at specified time points, indicated by arrows in A, and imaging was performed. Representative microscopy images are shown. **C,** population distribution of GFP fluorescence in the wild-type and (p)ppGpp^0^ strains at specified time points. **** denotes a two-tailed P-value, derived from a non-parametric Mann-Whitney test, P<0.0001. The image analysis is representative of three independent experiments.

Two possible causes of lower AbrB levels are reduced *abrB* transcription or AbrB protein synthesis. We therefore constructed a fusion of P*_abr_*to firefly luciferase (P*_abrB_*-luc) and monitored its activity, observing that P*_abr_* increased during approximately the same interval (∼200-250 min) as P*_hpf_* and P*_sigH_* (Fig. S7). Thus, changes in *abrB* transcription likely do not account for the observed decrease in AbrB abundance (Fig. 5B, C, ‘wildtype’). Alternatively, the decrease in AbrB abundance may be a consequence of the attenuation of global protein synthesis during the period of *hpf* activation (Fig. 2C, D). This attenuation is partially dependent on (p)ppGpp as a (p)ppGpp^0^ strain exhibited higher levels of OPP labeling during this interval (200-250 min) as compared to the wildtype parent (Fig. S8). To investigate if this was also true for AbrB-GFP, we compared its abundance in ppGpp^0^ and wildtype parent strains, observing that AbrB-GFP levels were higher in the ppGpp^0^ background (Fig. 5B, C, ‘(p)ppGpp^0^’). Thus, (p)ppGpp affects AbrB abundance, providing a plausible explanation for the dependence of increased expression of σ^H^ and HPF on (p)ppGpp.

The σ^H^ regulon is relatively large (30), suggesting that other genes could be regulated by (p)ppGpp similarly to *hpf*. To investigate this possibility, we employed scRNAseq (single cell RNA sequencing) (31) since FISH analysis of P*_hpf_*-GFP reporter indicates that *hpf* transcription in single cells is rather heterogeneous (Fig. 6A, B) We analyzed wildtype parent and (p)ppGpp^0^ strains at a time point near maximum P*_hpf_*-luc expression. The expression of numerous genes was significantly higher in the wildtype, including *hpf* (red) and *sigH* (blue) (Fig. 6C), consistent with observations reported here (Figs, 2, 4, respectively). Many of these genes are members of the σ^H^ regulon (green) including *rapC* (32), *kinA* (33), and *ftsZ* (34) (Fig. 6C). We generated firefly luciferase fusions to the promoters of several of these genes and explicitly assessed their dependence on (p)ppGpp. For example, expression of a P*_ftsAZ_*-luc fusion during post-exponential growth is dependent on (p)ppGpp (Fig. 7A). This dependence has physiological consequences as ppGpp^0^ cells are significantly longer than the wildtype parent (Fig. 7B, C), consistent with the observation that lower levels of FtsZ and FtsA attenuate cell division (35).

**Figure 6.**
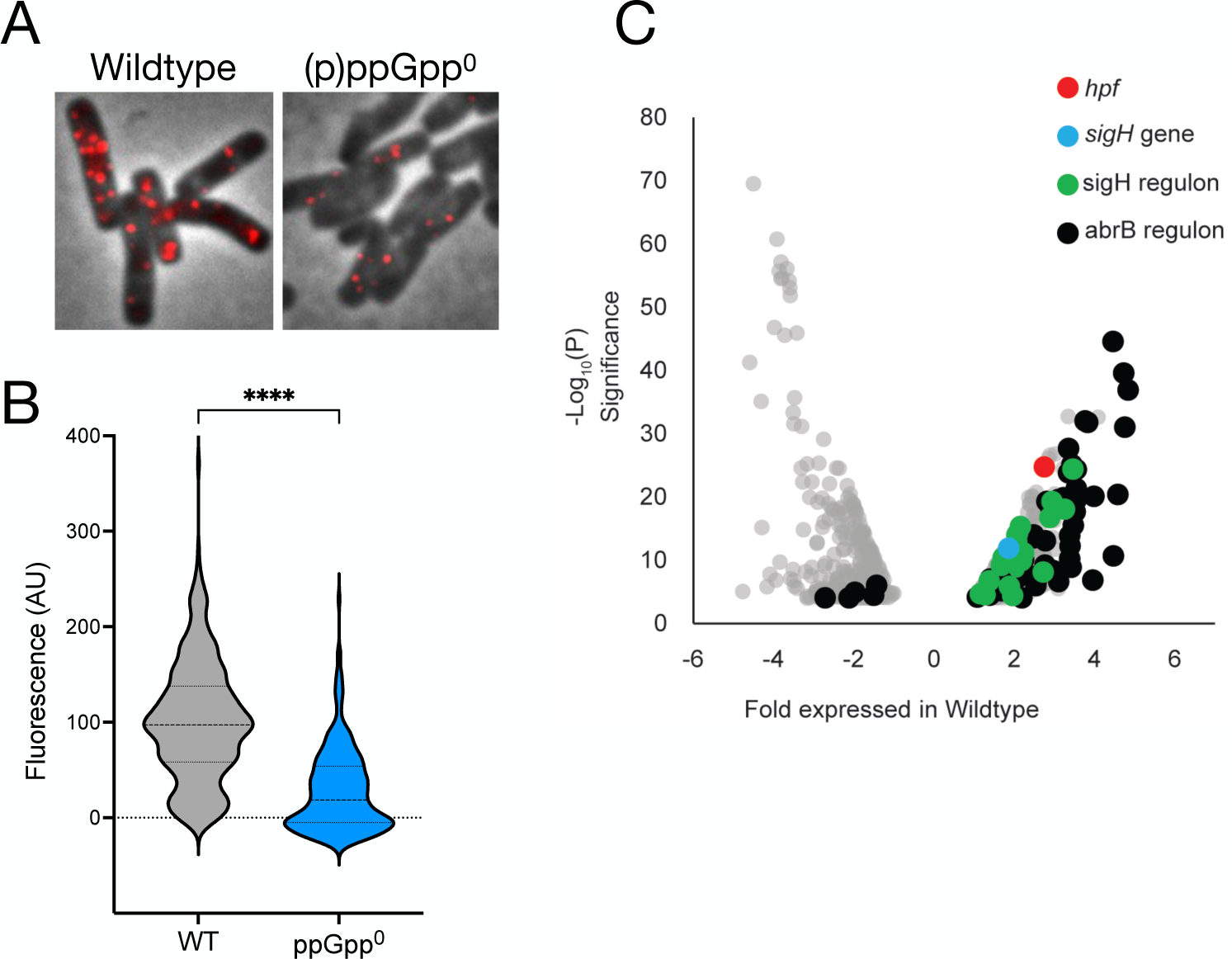
Genes and regulons upregulated in a (p)ppGpp^0^ dependent manner. **A,** aliquots of wildtype (JDB4450) and (p)ppGpp^0^ (JDB4828) strains expressing P*_hpf_*-GFP were collected at t∼240 min and analyzed by FISH using oligos complementary to *gfp*. **B,** population distribution of fluorescence. **** denotes a two-tailed P-value, derived from a non-parametric Mann-Whitney test, of P<0.0001. **C,** samples of wildtype (JDB4811) and (p)ppGpp^0^ (JDB4812) strains expressing P*_hpf_*-luc were collected at the time of maximal P*_hpf_* -luc (t=240 min) and analyzed using the scRNA-seq technique (31). The volcano plot denotes differential gene expression between the wild-type and a (p)ppGpp^0^ mutant, with each dot representing one gene. Indicated are *hpf* (red), *sigH* (blue), SigH regulon (green), AbrB regulon (black), and genes down-regulated in a (p)ppGpp^0^ mutant (grey). The fold differences in gene expression are indicated on the x-axis and the fold difference Log_10_(P) statistical significance is represented on the y-axis.

**Figure 7.**
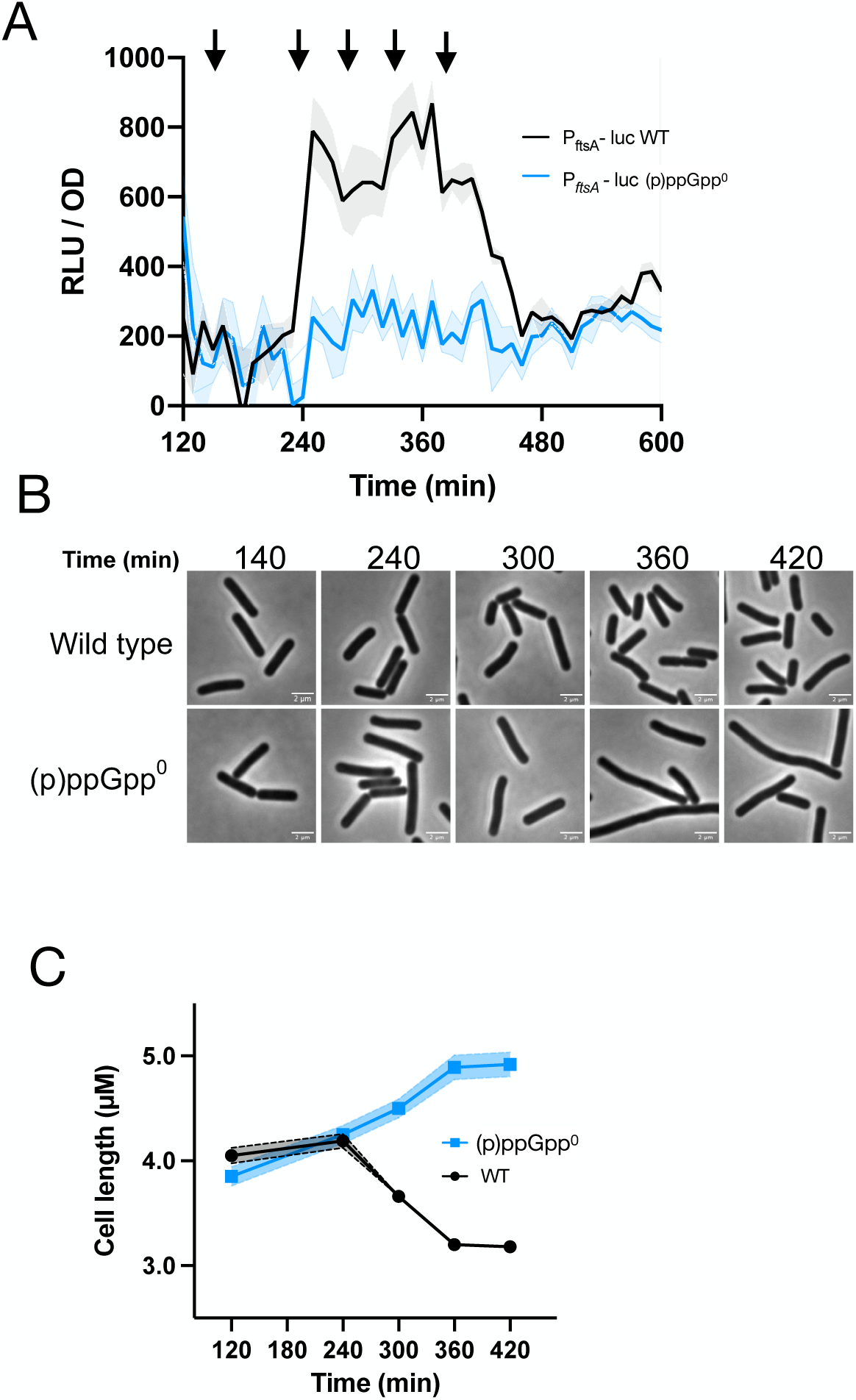
Role of (p)ppGpp in cell division. **A,** luminescence (RLU/OD_600_) of P*_ftsAZ_*-luc*-* containing wildtype (black, JDB4829) and (p)ppGpp^0^ (blue, JDB4830) strains; **B**, representative microscopy images of the wildtype and (p)ppGpp^0^ strains taken at time points indicated in A (arrows); **C,** average cell length (µM) measured over time during growth for the wildtype (black) and (p)ppGpp^0^ (blue) strains.

Expression of *rapF* is also σ^H^-dependent (32) and (p)ppGpp-dependent (Fig. 6). Consistently, a P*_rapF_*-luc fusion is dependent on (p)ppGpp, at least during post-exponential growth (Fig. 8A). P*_rapF_*drives expression of the genes encoding RapF and its inhibitor PhrF, and together both proteins control the transcription factor ComA (36), the master regulator of competence gene expression (37). We therefore investigated if the dependence of P*_rapF_* expression on (p)ppGpp has consequences for the development of competence. We obtained cells from both a wildtype parent and a (p)ppGpp^0^ strain at a time point associated with high P*_rapF_*expression (Fig. 8A, arrow). We incubated the cells with genomic DNA from a suitable donor strain, observing that the ppGpp^0^ strain was reduced >100-fold in transformability as compared to the wildtype parent (Table 1). A third σ^H^-dependent gene we identified as being positively regulated by (p)ppGpp is *spo0A*, encoding the master regulator of sporulation and other post-exponential processes. Although P*_spo0A_*-luc is expressed at a high level early in growth, it exhibits a (p)ppGpp-dependent increase in activity at a similar point in the growth curve (Fig. 8B) as P*_hpf_-*luc. Consistently, the production of heat-resistant spores is also dependent on (p)ppGpp in S7/glucose (Table 2). However, this effect was media-specific as production of spores in DSM was unaffected (Table 2).

**Figure 8.**
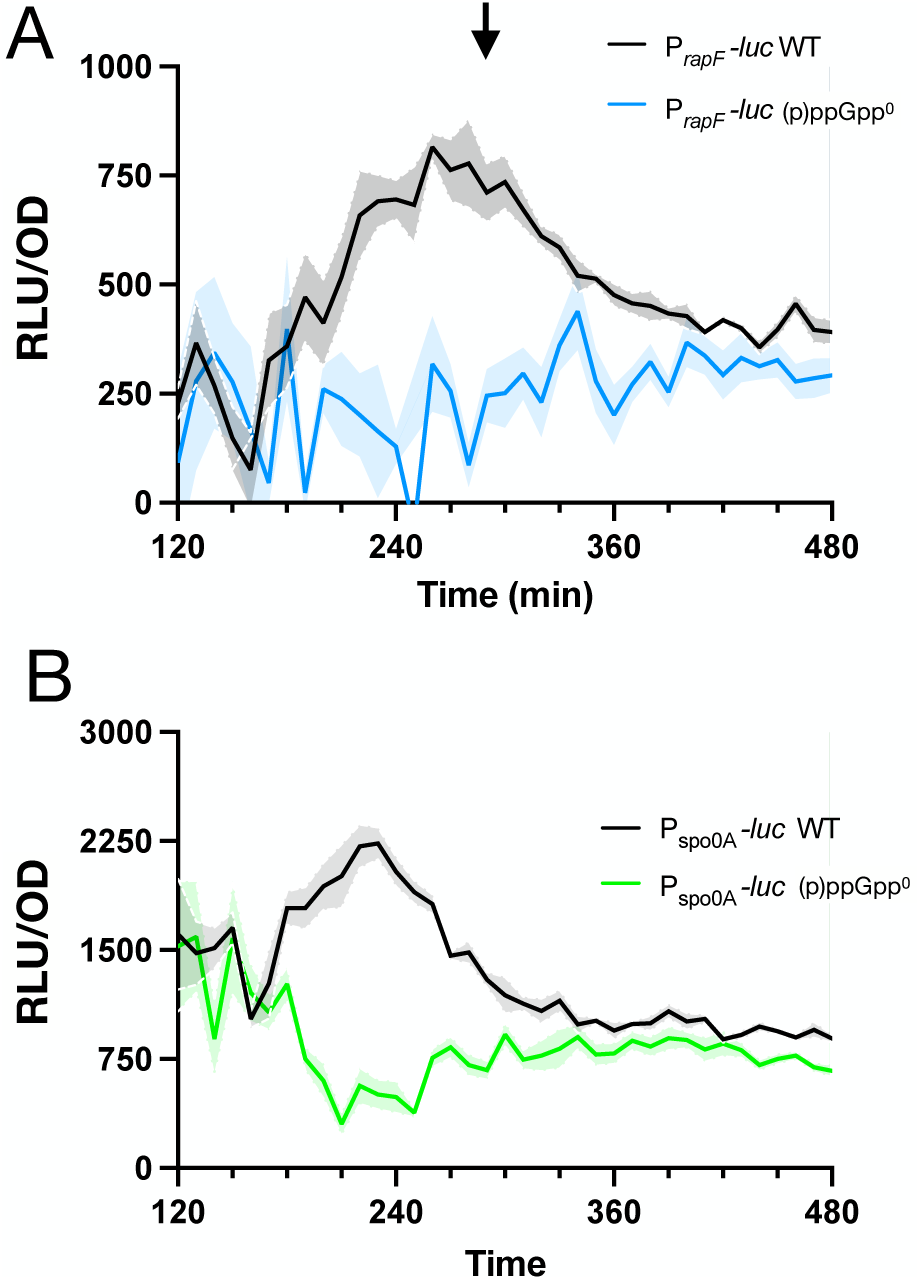
Role of (p)ppGpp in post-exponential gene expression. Luminescence (RLU/OD_600_) of wildtype (black) and (p)ppGpp^0^ (blue) strains containing **A,** P*_rapF_*-luc (JDB4831, JDB4832) **B,** P*_spo0A_*-luc (JDB4833, JDB4834). The arrow in A indicates the time when cells were collected for the competence assay.

**Table 1.** Transformability of wildtype and (p)ppGpp^0^ mutants.

| Genotype | Growth Media | CFU/ml | Transformants |
| --- | --- | --- | --- |
| Wild type | S7 | 1.64 X 10<sup>8</sup> | 233 |
| (p)ppGpp<sup>0</sup> | S7 | 0.90 X 10<sup>8</sup> | 0 |

Strains were grown in S7/glucose. Transformation was assessed by incubating 100 µl of culture with 1 µg genomic DNA (JDB4692), selecting for cm<sup>R</sup>.

**Table 2.** Sporulation efficiency of the WT and (p)ppGpp^0^ mutants under different nutrient conditions.

| Genotype | Growth Media | Viable/ml | Spores/ml | Sporulation efficiency |
| --- | --- | --- | --- | --- |
| Wild type | DSM | 2.10 x 10<sup>7</sup> | 1.93 x 10<sup>7</sup> | 92% |
| (p)ppGpp<sup>0</sup> | DSM | 2.55 x 10<sup>7</sup> | 2.21 x 10<sup>7</sup> | 87% |
| Wild type | S7 | 1.97 x 10<sup>7</sup> | .96 x 10<sup>7</sup> | 48% |
| (p)ppGpp<sup>0</sup> | S7 | 2.16 x 10<sup>7</sup> | 0.27 x 10<sup>3</sup> | .0013% |

Strains were grown under specified conditions and sporulation assessed after 48h. Viable/ml and Spores/ml refers to CFU/ml before and after heat treatment (80°C, 15 min), respectively. Efficiency is defined as the ratio of spores/ml to viable/ml. Shown is the result from a single experiment, similar results were obtained in three biological replicates.

## Discussion

These experiments demonstrate that post-exponential expression of the *hpf* gene encoding a ribosome hibernation factor is under control of the alarmone (p)ppGpp. Unlike direct mechanisms of transcriptional control by (p)ppGpp that involve its binding to RNA polymerase or to a specific transcription factor (38), *hpf* activation is indirect. (p)ppGpp attenuates global translation, and, as a consequence, the abundance of the transcriptional repressor AbrB falls, thereby increasing expression of its target gene, *sigH*, encoding σ^H^ required for the expression of *hpf.* σ^H^ is a central player in the transcriptional response of Gram-positive bacteria including *B. subtilis* to reduced nutrient availability (30). We demonstrate that (p)ppGpp participates in the σ^H^-dependent transcriptional changes that underly the post-exponential differentiation of *B. subtilis* into shorter cells that eventually become genetically competent or sporulate.

AbrB expression is also regulated by the Spo0A transcription factor. Spo0A inhibits *abrB* transcription, at least later in the transition phase when Spo0A is highly activated (Spo0A∼P)(39). However, since *abrB* transcription increases concomitantly with *sigH* activation (Fig. 4, Fig. S7), transcriptional repression by Spo0A is likely not involved in this phenomenon. Spo0A∼P also regulates the transcription of the AbrB inhibitor AbbA (40). However, deletion of *abbA* does not substantially affect *sigH* transcription (Fig. S9). Taken together, these data indicate that Spo0A is not important for the AbrB-dependent stimulation of *sigH* expression underlying *hpf* induction during early growth transition. In addition, scRNA-seq transcriptome analysis revealed that several genes of the *abrB* regulon are regulated in a ppGpp-dependent manner during the early growth transition (Fig. 6C). Thus, ppGpp inhibits *abrB* translation early during transition phase whereas Spo0A inhibits *abrB* transcription at later times (41).

(p)ppGpp reduces transcription of rRNA and ribosomal protein genes in Proteobacteria via direct interaction with RNAP and in Firmicutes by effects on GTP levels, as GTP is typically the initiating nucleotide of rRNA genes (8). However, the role of these mechanisms in gene *activation* dependent on (p)ppGpp (e.g., amino acid biosynthetic genes) remains unclear. Several models have been proposed including the affinity model where RNAP liberated from stable RNA synthesis results in an increase in free RNAP, facilitating increased transcription of amino acid biosynthetic genes (42). However, the data presented here suggests that activation can result from the reduction in protein synthesis mediated in part by the inhibition of translation by (p)ppGpp. Other mechanisms by which ppGpp could activate transcription include binding to a class of riboswitches found upstream of BCAA biosynthetic genes in some Firmicutes (43), but to our knowledge, this has not been demonstrated. (p)ppGpp binds the PurR transcriptional repressor of the *B. subtilis pur* operon, thereby increasing PurR DNA binding affinity and repression (44), suggesting that changes in DNA avidity following (p)ppGpp binding to a transcription factor could result in gene activation but this has not been demonstrated.

Although we have focused on investigating the effect of σ^H^ on *hpf* transcription, the *hpf* promoter also contains a binding site for σ^B^, the general stress transcription factor (16). A reporter containing a scrambled σ^B^ binding site (P*_hpf_*_-*sigB\**_*-*luc) also exhibits reduced *hpf* activity under our experimental conditions (Fig. S5). The effects of σ^B^ and σ^H^ are not overlapping since a strain expressing a reporter carrying both scrambled σ^B^ and scrambled σ^H^ binding sites (P_hpf-sigB*H*_-luc) exhibits essentially no *hpf* activity (Fig. S6). Thus, *hpf* appears to be an exception to the fact that σ^B^ and σ^H^ have mostly separate regulons (45). σ^B^ is typically thought of as a stress response (e.g., heat, ethanol, or salt shocks (46)) so it is not clear how the experimental conditions utilized in the present study fit this paradigm. However, σ^B^ is responsive to glucose exhaustion (47) and, intriguingly, to blue light (48, 49), both of which are stimuli that may be present during the growth conditions examined here.

The connection between (p)ppGpp and division has long been a subject of study, with the first observations of cell filamentation of an *E. coli* RelA mutant dating back >40 years (50). RelA overexpression leads to FtsZ overproduction suggesting a regulatory connection between (p)ppGpp and FtsZ (51). However, in contrast to the experiments described here (Figs. 6, 7), numerous transcriptomic experiments did not observe an effect of (p)ppGpp on *ftsZ* expression (e.g., (3, 4)), possibly because these studies used *E. coli* and experimental conditions different from those in the present study. Of note, a recent study observed (p)ppGpp, together with DksA, regulates division under basal conditions through downstream effects on transcription, although the mechanism was not identified (52). In *B. subtilis*, increased *ftsZ* transcription during early sporulation requires a σ^H^-dependent binding site in the *ftsAZ* promoter (34), suggesting that the increased expression we observed, concomitant with increased *spo0A* expression (Fig. 6A), could be related to sporulation, specifically to the formation of the asymmetric septum (53).

The requirement of (p)ppGpp for efficient sporulation in S7/glucose (Table 2) is consistent with previous reports (20) using DSM, a medium optimized for sporulation (54). Similarly, the defect in competence of a ppGpp^0^ strain (Table 1) is also consistent with earlier reports that used a defined competence medium (55). Thus, the (p)ppGpp dependence is growth medium-independent, suggesting that an intrinsic aspect of *B. subtilis* post-exponential physiology results in the well characterized differentiation into distinct cell fates (56). While the origins of this diversity remain unclear, single cell variation in (p)ppGpp abundance could play a role given that expression of genes encoding the regulatory networks underlying these differentiation pathways is dependent on (p)ppGpp (Figs. 6C, 7A, 8A, B). Consistently, expression of *hpf* - one of those genes - exhibits significant single cell variability (Fig. 6A, B).

In summary, our findings demonstrate extensive transcriptional reprogramming orchestrated by (p)ppGpp during the reduction in the growth rate that follows exponential growth. This mechanism therefore couples general nutrient availability, reflected in amino acid abundance sensed by Rel, with differential gene expression. Importantly, this anticipatory mechanism, likely initiated by the (p)ppGpp-dependent translation attenuation of the AbrB repressor, facilitates the emergence of potentially energetically costly adaptations before the cells enter a starvation regime where such changes would be much more difficult to implement.

## Acknowledgements

We acknowledge helpful advice from members of our laboratories. This work was supported by NIH R35GM141953 to JD and Startup funds from the UNC Department of Microbiology and Immunology to AR.

## Methods

### Strains

Strains, plasmids, and primers used are listed in Table S1, S2, S3, respectively. All strains are derivatives of *B. subtilis* 168 unless otherwise noted. To construct the P*_hpf_*-luciferase fusion (P*_hpf_-* luc), the region between -15 and -146 nucleotides upstream of the *hpf* start codon was amplified from JBD1772 genomic DNA. The pSD47 plasmid harboring the firefly luciferase gene was used as the vector backbone and amplified with compatible overhangs (19). This plasmid contains 3’ and 5’ flanking regions homologous to the *B. subtilis sacA* loci. PCR-purified P_hpf_ was cloned upstream of the firefly luciferase gene using Gibson assembly master mix (NEB) to create pSN09 that was transformed into *B. subtilis* 168 (JDB1772), selected for cm^R^, resulting in the P*_hpf_-*luc reporter at *sacA (*JDB4811). To construct various (p)ppGpp synthase mutants, gDNA isolated from the P_hpf_-luc reporter strain JDB4811 was transformed into *relA^Y308A*^* selecting for cm^R^ making JDB4816. To create a (p)ppGpp^0^ mutant, JDB4816 was made competent and transformed with gDNA of JBD3652 (20). The transformants were then sequentially selected for kan^R^ followed by tet^R^ to construct JDB4812. A similar approach was utilized to construct P*_sigH_,* P*_abrB_* and P*_ftsAZ_* luciferase reporter fusions in the wildtype and (p)ppGpp^0^ strains. Next, the consensus binding sites for the regulatory factors were scrambled to create P*_hpf-codY*,_* P*_hpf-sigB*,_* P*_hpf-sigH*,_ and* P*_hpf-sigB*H*._* These scrambled mutants were synthesized using IDT gene blocs, with overhangs that facilitate cloning the promoter mutants upstream of the firefly luciferase reporter using the Gibson technique. The strains harboring promoter mutants were constructed by transforming these plasmids into JBD1772. For FISH, P*_hpf_* promoter was cloned upstream of GFP^mut2^ fluorescent protein in pAF51 plasmid, which has the 3’ and 5’ flanking regions homologous to the *pyrD* loci in *B. subtilis* genome. The construct was then transformed into JDB1772, making JDB4450, followed by the introduction of (p)ppGpp synthase mutations to make JDB4828.

### Media, growth conditions and luminescence assay

Bacteria were grown in S7_50_ minimal media comprising 1X MOPS buffer (Teknova M2101) diluted in reagent grade water (Teknova W0225), supplemented with 1% glucose, 0.1% glutamic acid, 0.01% Casamino acids, 50mM NaCl, 40µg ml^-1^ tryptophan and 132 mM potassium phosphate buffer (Teknova M2102). The 10X S7_50_ salt solution is prepared by dissolving 50mM MOPS, 10mM (NH_4_)_2_SO_4_, and 5mM KH_2_PO_4_ in ddH_2_0, buffered to pH 7.0 with 5M KOH, filter sterilized and stored at 4℃. The 100X trace metal solution (Teknova 2M2755) contains 200mM MgCl_2_, 70mM CaCl_2_, 5mM MnCl_2_, 0.1mM ZnCl_2_, 100µg ml^-1^ thiamine HCl, 0.5mM FeCl_3_, & 2mM HCl. For uracil auxotroph *pyrD* mutants, 40 µg ml^-1^ uracil was supplemented.

Cultures of bacterial strains from a single colony were grown in S7_50_ minimal media until cultures reached an OD_600_ = 0.3-0.4. The cultures were diluted to an initial OD_600_ = 0.05 in S7_50_ minimal media supplemented with 4.7 mM D-luciferin (Goldbio) in 96-well flat bottom white-sided plates (Greiner Bio-One 655098) and grown at 37 °C with continuous shaking in a Tecan Infinite 200 plate reader. Measurements of luminescence and OD_600_ were taken at 10 min intervals. Media only and luciferin controls were used for background subtraction for OD_600_ and luminescence, respectively. All cultures were grown in triplicate. The growth and RLU/OD were plotted against time on the X-axis and RLU/OD on the Y-axis.

### Immunoblot analysis

Based on the P*_hpf_*-luc activity, at specified time intervals, 2 ml aliquots of cells were collected, resuspended in 100 µl Lysis buffer (20 mM Tris-Cl pH 8.0, 100 mM NaCl, 1:100 diluted protease inhibitor cocktail (P8340, Sigma), 0.1 mg ml^-1^ lysozyme, 10mg ml^-1^ DNase, 100 µg ml^-1^ RNase) and incubated at 37℃ for 15 minutes. Protein concentration was determined with a Coomassie plus Bradford detection kit (BioRad). An equal protein concentration from each sample was loaded onto a 15% polyacrylamide gel. The steady-state levels of HPF and SigA were analyzed using anti-HPF (1:10000) and anti-SigA primary antibodies (1:10000), respectively. The primary antibodies were detected using horseradish peroxidase-conjugated goat, anti-rabbit immunoglobulin G (BioRad) and the western detection kit as described by the manufacturer (PerkinElmer). The anti-SigA antibody was a kind gift from Niels Bradshaw (Brandeis University) and the anti-HPF antibody was previously generated in our laboratory (22).

### Microscopy and Image Analysis

Microscopy was performed on live or fixed cells immobilized on 1% agarose prepared with S7_50_ media. Imaging was performed using a Nikon 90i or a TE2000 microscope with a Phase contrast objective (CFI Plan Apo Lambda DM ×100 Oil, NA 1.45), an X-Cite light source, a Hamamatsu Orca ER-AG, and the following filter cubes: FITC and mCherry. Phase contrast and fluorescence images (GFPmut2/mCherry) of bacterial cells immobilized on agarose pads were acquired. The image stacks were analyzed using Fiji with the help of the MicrobeJ plugin (57). The straighten and intensity options in the MicrobeJ plugin were used to measure the average fluorescence per pixel within each cell. Mann-Whitney statistical test was performed using Prism to ascertain the significance of the fold differences between the strains. A non-fluorescent control strain was used to subtract background and autofluorescence in each channel.

### OPP Labelling

OPP (O-propargyl-puromycin) labelling was performed using the Click-iT™ Plus OPP Alexa Fluor™ 594 Protein Synthesis Assay Kit (Invitrogen) as described (19). Briefly, P_hpf_ luciferase reporter activity was monitored, and at the specified time intervals, 0.6 ml cells were collected and transferred to disposable glass tubes. OPP was added to a final concentration of 13 µM, and the tubes were incubated at 37℃ for 15 min. Cells were harvested by centrifugation at 10000 RPM for 1 min and resuspended in 100 µl of PBS containing 3.7% formaldehyde for fixation. Cells were fixed for 10 min, harvested and permeabilized with 100µl of 0.5% Triton-X 100 in PBS for 15 min. Cells were conjugated with the fluorophore Alexa Fluor™ 594 using 100µl of 1X Click-iT cocktail for 20 min in the dark following the manufacturer’s guidelines. Cells were harvested and washed once using Click-iT rinse buffer and then re-suspended in 20-40 µl of PBS for microscopy. Phase contrast and fluorescence images (mCherry) of bacterial cells immobilized on agarose pads were acquired.

### GFPmut2 probes and Fluorescent *in situ* hybridization

The GFPmut2 FISH probes were designed and synthesized with CAL Fluor® Red 590 Dye (LGC Biosearch Technologies). FISH imaging was performed as described (31) with slight modifications. Briefly, 2 ml cells from mid or late-log cultures grown in S7_50_ minimal media were collected and fixed with 1% formaldehyde (final concentration) at room temperature for 30 min. The cells were then harvested by centrifugation at 6000 RPM for 3 min at room temperature and washed with 0.02% saline sodium citrate (SSC, Invitrogen). The cell pellets were resuspended in 300 µl MAAM mix (4:1 V:V dilution of methanol to glacial acetic acid) and incubated at -20 ℃ for 15 minutes, followed by 1X PBS (from 10X PBS, Invitrogen) wash to remove traces of MAAM. Cells were permeabilized in 200µl PBS containing 350 U µl^-1^ of lysozyme (Epicenter ready-lyse) for 30 min at 37°C. After permeabilization, cells were washed once with 500 µl PBS. The cells were resuspended in 100 µl Stellaris® RNA FISH Hybridization Buffer containing 10% formamide and 12.5 µM reconstituted GFP^mut2^ oligo probes for hybridization. The cell-probe mix was incubated in a 30°C water bath overnight. Cells were harvested and washed with 500 µl of reconstituted Stellaris® RNA FISH Wash Buffer A containing 10% formamide and incubated at 30°C water bath for 1 hour. The washing step was repeated one more time to remove the excess probe. The cells were harvested and resuspended in 0.5 ml Stellaris® RNA FISH Wash Buffer B and incubated for 5 minutes at room temperature. Finally, the cells were harvested and resuspended in 20-40 µl of PBS with 1µl SlowFade™ Gold Antifade Mountant (Invitrogen). A non-fluorescent control strain, treated with a GFP^mut2^ probe, was used as a control to subtract background and autofluorescence in each channel. Phase contrast and fluorescence images (mCherry) of bacterial cells immobilized on agarose pads were acquired.

### Transcriptomic analysis using ProBac-seq microfluidic encapsulation of wildtype and ppGpp^0^ cells

At the indicated timepoints wildtype and ppGpp^0^ cells were fixed in 1% formaldehyde at room temperature for 30 min. Fixed cells were pelleted by centrifugation at 6000 RPM for 3 min at room temperature and washed with 0.02% saline sodium citrate (SSC, Invitrogen). The cell were pelleted again and resuspended in 300 µl MAAM mix (4:1 V:V dilution of methanol to glacial acetic acid) and stored at 20 °C until processing. ProBac-seq sample processing was done as described previously (31). Briefly, samples were pelleted and resuspended in PBS to remove MAAM. Following PBS wash to remove MAAM the cells were incubated with lysozyme (Readylyse, Epicenter) for 30 min at room temperature for cell wall hydrolysis. After an additional pelleting step and PBS wash the samples were further permeabilized by a 5 min incubation in PBS-Tween (PBS with 1% final concentration of Tween-20 detergent) and a subsequent wash in PBS to remove the detergent. Permeabilized cells were resuspended in 100 μl of probe binding buffer (5 × SSC, 30% formamide, 9 mM citric acid (pH 6.0), 0.1% Tween 20, 50 μg ml−1 heparin and 10% low molecular weight dextran sulfate purchased from Molecular Instruments as Probe Binding Buffer – Cells in Suspension). A transcriptome-wide probe-set containing approximately 30,000 mRNA binding probes designed against the *B. subtilis* genome (31) was added to each sample before incubate the samples overnight at 50^°^C. After overnight incubation unbound probes were removed by a series of seven washes with probe wash buffer (5 × SSC, 30% formamide, 9 mM citric acid pH 6.0, 0.1% Tween 20 and 50 μg ml-1 heparin (Molecular Instruments)) and 2 final washes in PBS. Probed and washed cells were enumerated using a flow-cytometer and ∼10,000 cells per condition were processed using the ProBac-Seq protocols as described (31). In short, 51 μl cell suspensions in PBS were mixed with 19 μl ddPCR master mix (Biorad), 3 μl in-droplet PCR primer, 2 μl reducing agent (10X genomics) to provide a 75μl reaction mix. 70 μl of this reaction mixture was loaded onto 10X Genomics microfluidic chip-G devices alongside 10X barcoding beads and emulsion oil. The devices were run using a 10X chromium controller and emulsions were processed to produce barcoded libraries as described previously. Libraries were sequenced using an Illumina nextSeq2000 100-cycle kit and fastQ-files were demultiplexed, mapped and normalized using the 10X cellRanger and Seurat software packages using the standard analysis pipelines and default parameters. To compare gene expression between wildtype and ppGpp^0^ strains, we grouped all cells from a condition and compared the normalized signal across conditions using duplicate samples. Fold changes and P-values calculated in the standard cellRanger differential gene expression pipeline (58) were used to produce volcano plots and genes annotated to be a part of the SigH and AbrB regulons in Subtiwiki (59) were highlighted.

